# Predicting the risk of sarcopenia in elderly patients with patellar fracture: Development and assessment of a new predictive nomogram

**DOI:** 10.1101/702548

**Authors:** Chen Yisheng, Tao Jie

## Abstract

**Purpose:** This study was aimed at developing a risk prediction model for postoperative dysplasia in elderly patients with patellar fractures in China.

**Patients and methods:** We conducted a community survey of patients aged ≥55 years who underwent surgery for patellar fractures between January 2013 and October 2018, through telephone interviews, community visits, and outpatient follow-up. We established a predictive model for assessing the risk of sarcopenia after patellar fractures. We developed the prediction model by combining multivariate logistic regression analysis with the least absolute shrinkage model and selection operator regression (Lasso analysis). The predictive quality and clinical utility of the predictive model were determined using C-index, calibration plots, and decision curve analysis. We conducted internal sampling methods for qualitative assessment.

**Result:** We recruited 61 participants (males: 20, mean age: 68.1 years). Various risk factors were assessed, and low body mass index and diabetes mellitus were identified as the most important risk factors (P<0.05). The model showed a good prediction rate (C-index: 0.909; 95% confidence interval: 0.81–1.00) and good correction effect. The C-index remained high (0.828) even after internal sample verification. Decision curve analysis showed that the risk of sarcopenia was 8.3–80.0%, suggesting good clinical practicability.

**Conclusion:** Our prediction model shows promise as a cost-effective tool for predicting the risk of postoperative sarcopenia in elderly patients based on the following: advanced age, low body mass index, diabetes, longer postoperative hospital stay, no higher education, no postoperative rehabilitation, removal of internal fixation, and less outdoor exercise.

## Introduction

In 2010, the European Working Group on Sarcopenia in Older People (EWGSOP) and the International Symposium on Sarcopenia in 2011 established a unified definition of sarcopenia, which highlighted that sarcopenia occurs as a result of a gradual decline in muscle mass. Sarcopenia is characterized by a gradual and generalized loss of the strength and function of skeletal muscles.^1,2^ It has be closely associated with diabetes as well as the risk of subsequent falls, fractures, and physical disabilities.^2-6^ Therefore, the prevention of sarcopenia can help reduce the risk of falls and secondary fractures, thereby improving the overall prognosis of patients with fractures.^7,8^

The occurrence of fractures is an important risk factor for sarcopenia.^9^ Patellar fracture is a common fracture of the lower limb ^10^ and may entail long-term bed rest for patients, which is an important cause of sarcopenia.^11^ The patella is an important component of the knee joint, playing an important role in the attachment of the quadriceps muscle to the patellar ligament and thereby maintaining the stability and functioning of the knee joint.

However, the pathogenesis of sarcopenia is multi-centric involving various factors, including those related to disease (such as health status and work intensity), treatment (such as surgery and rehabilitation programs and other medical-related issues), and patients (such as age, body mass index [BMI], gender, and education level). In addition, ageing and diabetes have also been implicated as important risk factors for sarcopenia.^12-14^ The risk of sarcopenia is reported to be high among patients with diabetes.^15^ Decrease in muscle mass and the resulting abnormal gait and slow walking speed has been reported as the mechanism underlying sarcopenia.^16^

Although several variables that affect sarcopenia have been reported, no method has yet been identified for systematic assessment to predict the risk of post-fracture muscle reduction; therefore, it is necessary to develop accurate predictive tools and early individualized interventions to prevent muscle loss. In 2014, the Asian Working Group for Sarcopenia (AWGS) established effective criteria for the diagnosis of sarcopenia based on the characteristics of the Asian population; these criteria are being widely used to assess the occurrence and risk factors of sarcopenia.^17^ We believe that the AWGS criteria can be used to predict the occurrence of sarcopenia after patellar surgery by establishing an effective nomogram model.

The aim of this study was to develop an effective and simple tool for predicting sarcopenia in patients undergoing post-fracture patellar surgery. Only the data available for period after the patient’s hospitalization for fracture were used to evaluate the risk of sarcopenia in patients with patella fracture in order to study the effects of early prevention of sarcopenia and obtain accurate guidance for clinical work.

## Patients and methods

### Patients

This study was designed as a retrospective investigation of the prognosis of elderly patients with patella fracture. The study protocol was approved by the ethics committee of the First People’s Hospital affiliated to Shanghai Jiao Tong University Medical School (approval no. 2019SQ059). The study was performed in accordance with the stipulations of the Helsinki declaration. We first identified patients residing in Shanghai, China, who underwent patellar surgery at the Shanghai First People’s Hospital between January 2013 and October 2018. Between February 2019 and June 2019, we conducted a questionnaire-based survey through appointment-based telephonic interview, outpatient services, and community follow-up. Data for all the patients were collected by the same researcher. Written informed consent was sought from all enrolled subjects before participation in the study; all participating patients provided consent. In this study, we included only fracture patients aged ≥55 years, since postoperative sarcopenia after fractures is rare in young patients. The laboratory blood test results taken into consideration were those obtained for the preoperative fasting samples of venous blood.

### Inclusion criteria and exclusion criteria

The inclusion criteria for the survey were as follows: (1) diagnosis of sarcopenia according to the AWGS criteria, (2) age ≥55 years, (3) availability of complete data for baseline clinical characteristics (age, body mass index, etc.) and follow-ups, (4) patients had been managed with self-care before surgery (5) basic communication skills.

Patients were excluded from the study if they had major illnesses, serum creatinine levels of >2.5 mg/dL, liver enzyme levels elevated to more than twice the upper limit of the normal within 6 months prior to the start of the study, unexplained fever or infection, acute heart failure, or organ failure such as renal failure. Patients were also excluded if they did not cooperate with the completion of the basic diagnostic tests. Patients who developed life-threatening conditions during follow-up or conditions that could interfere with the interpretation of the study results were also excluded.

Seventy-three patients were screened, of which 61 were found to be eligible according to the inclusion criteria and availability of the completed questionnaire. The patients included in the final analysis included 20 males (age: 57–89 years, mean age: 67.6 ± 8.1years) and 41 females (age: 59–90 years, average age: 68.3 ± 7.7 years). These 61 patients were included in the final analysis.

## Methods of assessment

The diagnostic criteria established by the 2014 Asian Working Group for Sarcopenia (AWGS) defines sarcopenia by a walking speed of ≤0.8 m/s and measuring muscle mass (BIA) of ≤7.0 kg/m^2^ for males and ≤5.7 kg/m^2^ for females.^17^ The pace of decline in walking speed is closely related to the prognosis of patients.^18,19^ Hence, we used a fixed distance of 6 m, as recommended by AWGS, to measure the subject’s daily walking speed. For patients with a walking speed of ≤0.8 m/s, we used the bioelectrical impedance analysis (BIA) to assess muscle mass. The results of BIA are very similar to those of double-energy X-ray absorptiometry and magnetic resonance imaging; BIA also offers the advantages of safety, technical simplicity, low cost, and high patient compliance. ^20,21^ Muscle mass and BMI of the extremities were measured by using a bioimpedance meter (TANITA RD-953, Japan).

### Medical history and basic data collection

The questionnaire contained questions pertaining to various domains, including history of hypertension, diabetes, osteoporosis, malignant tumors, myocardial infarction, and falls as well as educational level, income, and duration of exercise. With regard to smoking habit, patients were classified as current smokers and non-smokers. Alcohol intake was calculated according to the average weekly alcohol consumption. The cut-off for exercise duration was 2 h/week, with duration of <2 h/week being considered as reduced duration. All patients had medical insurance that is provided for Shanghai residents. Additionally, we also reviewed the patients’ medical history and collected their laboratory blood test results.

### Statistical analysis

Statistical analysis was performed using R software (version 3.5.2). All the pre-determined factors described above were included in the LASSO analysis to reduce the dimensionality of the data, and the best predictors were identified (odds ratio of the OR values of these characteristic factors have 95% confidence intervals and P values <0.05).^22-24^ The multivariate logistic model was then used to analyze the factors selected with the LASSO regression model to establish a prediction model.^23^ The collected patient data can be used to generate a predictive model that accounts for all the best predictors of sarcopenia, which can be used for the prognostic evaluation of patients with patellar fractures.^25,26^ Next, we plotted the calibration curve to assess the accuracy of the nomogram. However, simple evaluation with the calibration curve would be incomplete.^27^ In addition, we also calculated the C-index and plotted the area under curve (AUC) curve to quantify the predictive power of the nomogram. We then performed further iterations (10,000 resampling) on the nomogram using the R language package for a more accurate measurement of C-index.^28^ Finally, we used the decision curve model to determine the net rate of return in order to test the clinical utility of the nomogram.^29^ The net rate of return was calculated by subtracting the number of all false-positive patients from the number of true-positive patients and comparing the negative effects of no intervention with those of excessive intervention.^30^

## Results

### Basic characteristics of the data

We obtained the completed questionnaires from 61 patients (20 males, 41 females; mean age 68.1 years) with patellar fractures. Patients were classified as those with (19 patients) or without sarcopenia (42 patients), as according to the diagnostic criteria established by the AWGS. Details of the patient characteristics are provided in Tables 1 and 2.

**Table 1:**
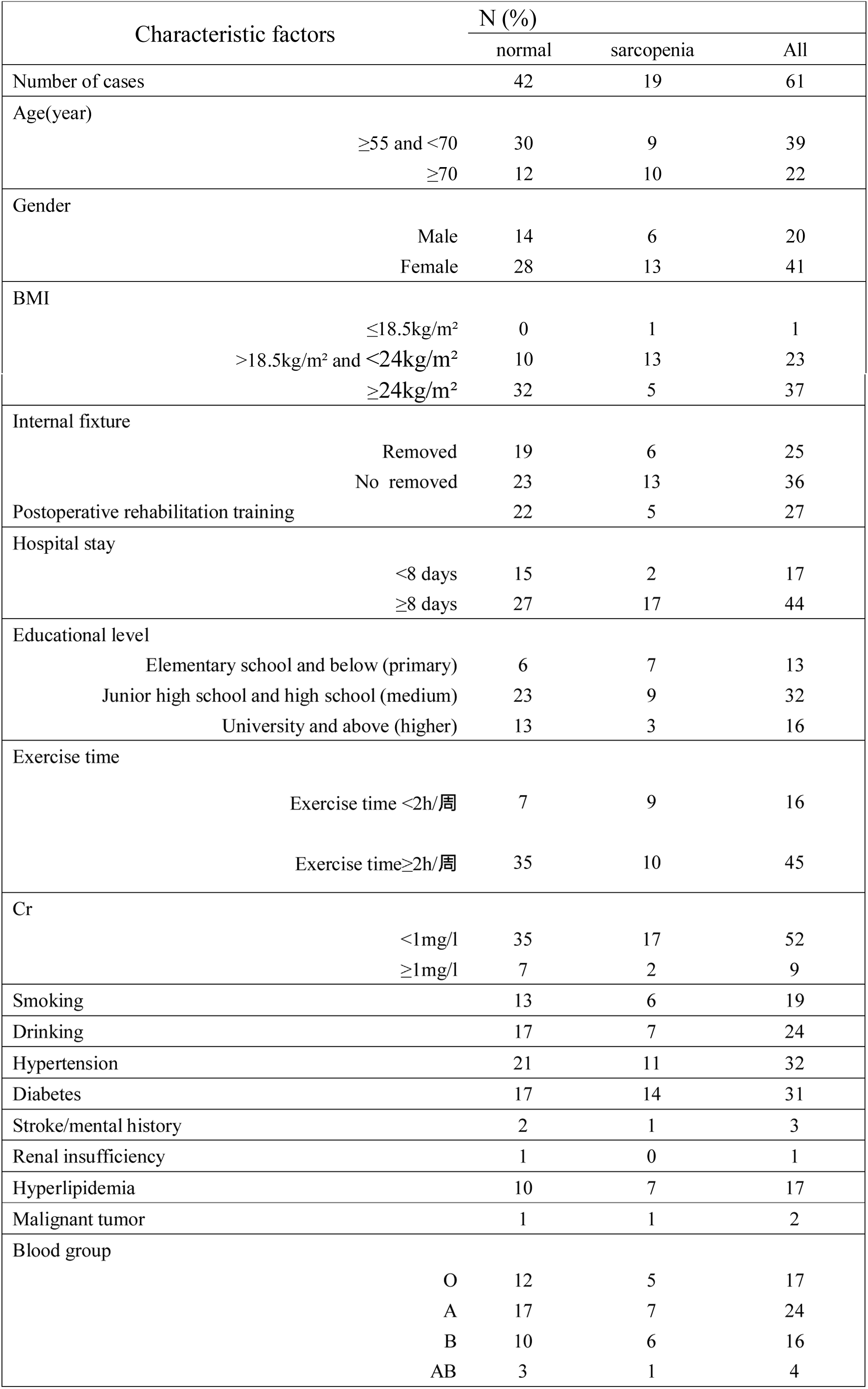
Relationship between sarcopenia and associated risk factors in elderly patients with patella fractures.

**Table 2:** Relationship between sarcopenia and associated risk factors in elderly patients with patella fractures.

|  | Normal | Sarcopenia | P level |
| --- | --- | --- | --- |
| Age ( year ) | 66.9±7.2 | 70.7±8.4 | 0.10 |
| Weight (kg) | 63.1±9.4 | 57.6±7.0 | <0.05 |
| Height (cm) | 157±10 | 161±13 | 0.25 |
| BMI(kg/m<sup>2</sup>) | 25.7±2.8 | 22.4±2.5 | <0.005 |
| Cr(mg/l) | 0.80±0.19 | 0.70±0.19 | 0.32 |

**Table 3:**
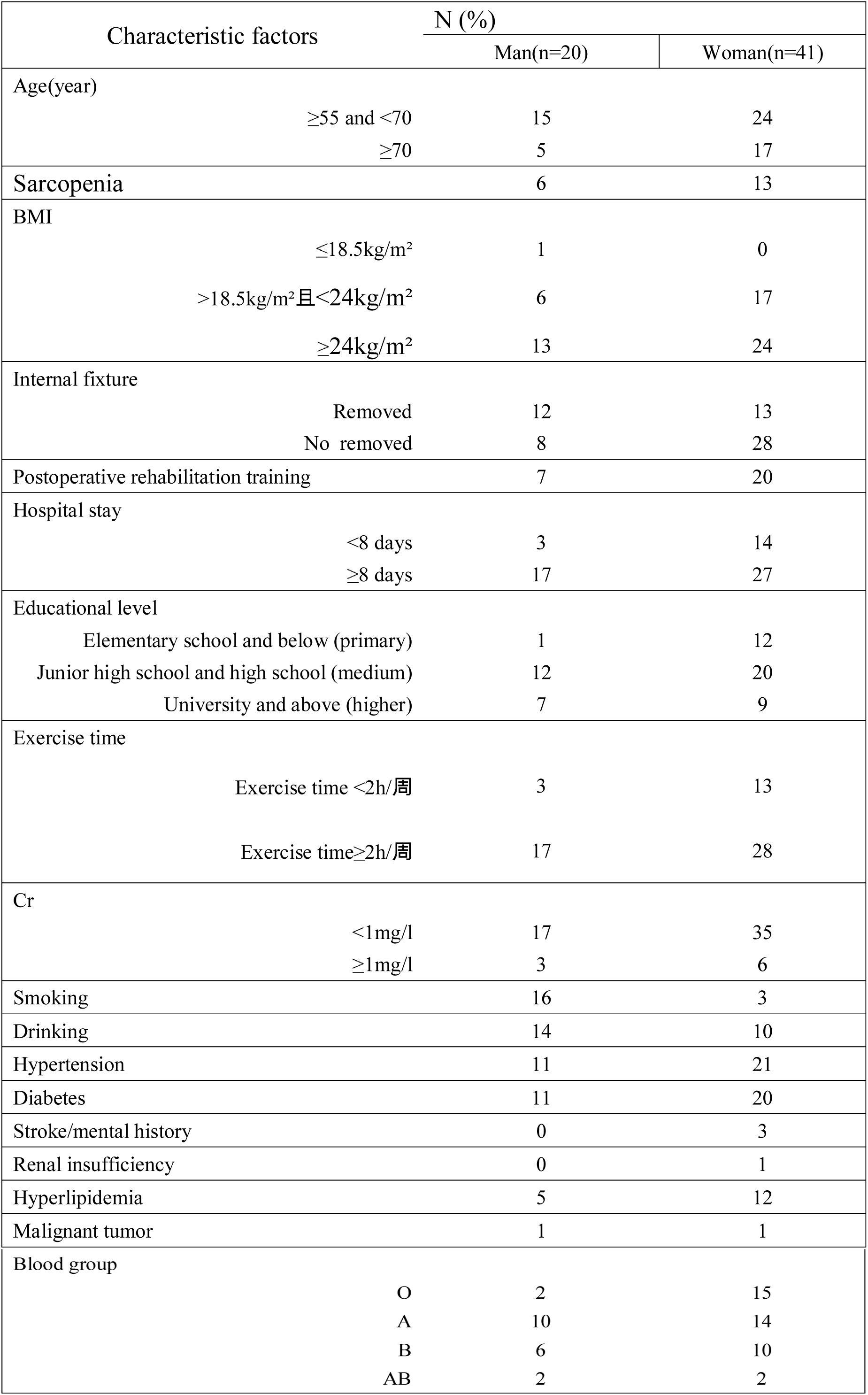
Comparison between factors of sarcopenia in man and woman.

### Screening of predictors

We used the LASSO regression model (Figures 1a and b) to decrease the factors in this study from 18 to 8 major parameters, which were age, BMI, education level, postoperative rehabilitation, diabetes, postoperative hospital stay, removal of internal fixation, and regular outdoor exercise.

**Figure 1.**
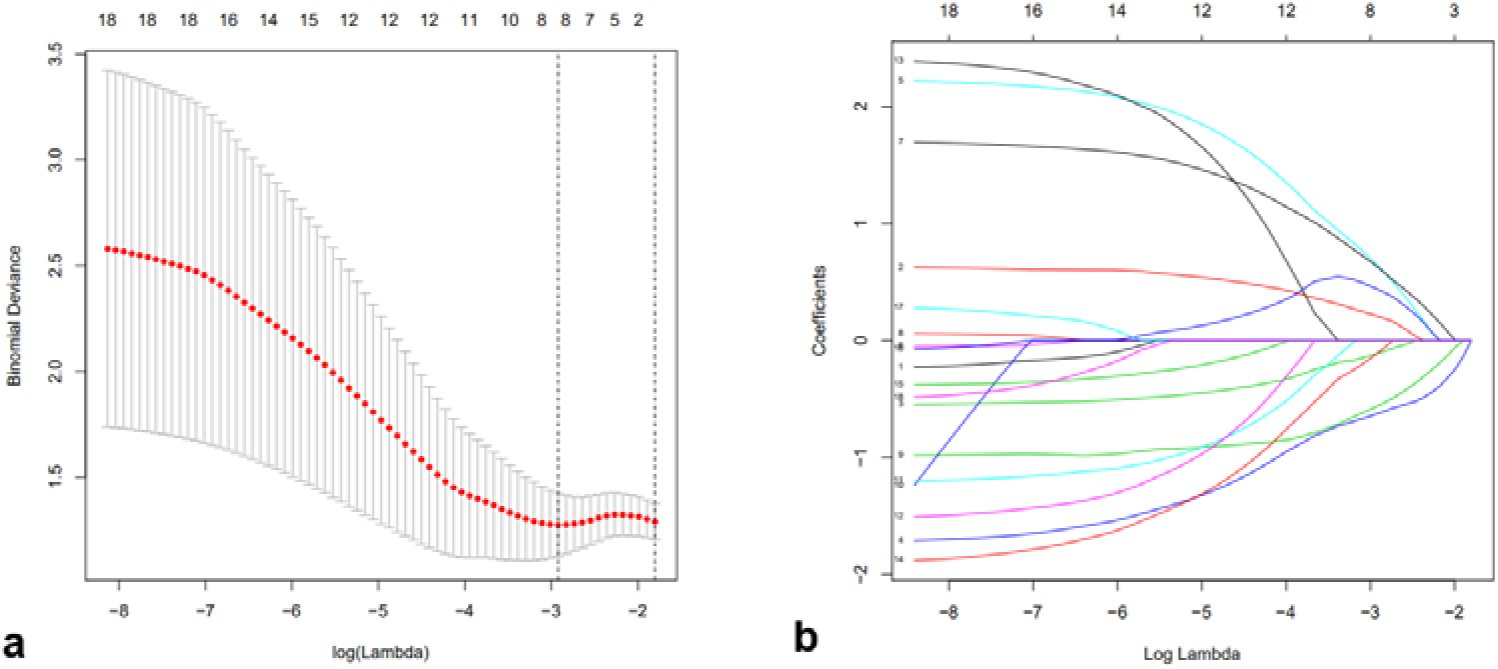
a. In the Lasso model, the choice of the optimal parameters used a five-fold cross-validation approach. Using the partial likelihood anomaly curve and the log (lambda) plot, the vertical line is drawn at the optimal value to obtain the included feature factors. b. The lambda curve generates a profile based on the log (lambda) sequence. Vertical lines were drawn at the values selected by the five-fold cross-validation method, and eight characteristic factors were selected.

### Construction of a personalized prediction model

Using the eight abovementioned predictors identified with the logistic model analysis (Table 2), we used the RMS package of R software to construct the column diagram, as shown in Figure 2 (see table 4 for details, p<0.05).

**Table 4.** Characteristic factors of predictive models of postoperative sarcopenia in elderly patients.

| Intercept and variable | Prediction model |  |  |
| --- | --- | --- | --- |
| | $\beta$ | Odds ratio (95%) | P-value |
| Intercept | -0.9156324 | 0.400(0.010-11.200) | 0.60 |
| age≥70s | 0.8087648 | 2.245(0.393-13.537) | 0.36 |
| BMI≥24kg/m<sup>2</sup> | -2.0199635 | 0.130(0.019-0.712) | 0.0251* |
| education level | -0.2667654 | 0.766(0.209-2.748) | 0.68 |
| No rehabilitation training | 1.4917357 | 4.445(0.842-31.395) | 0.0964 |
| diabetes | 2.1402414 | 8.501(1.501-71.919) | 0.0257* |
| exercise | -1.0614159 | 0.346(0.047-2.290) | 0.27 |
| internal fixation removal | -1.6419964 | 0.194(0.021-1.215) | 0.10 |
| length of hospital stay | 0.2588113 | 1.300(0.136-15.090) | 0.82 |

**Figure 2.**
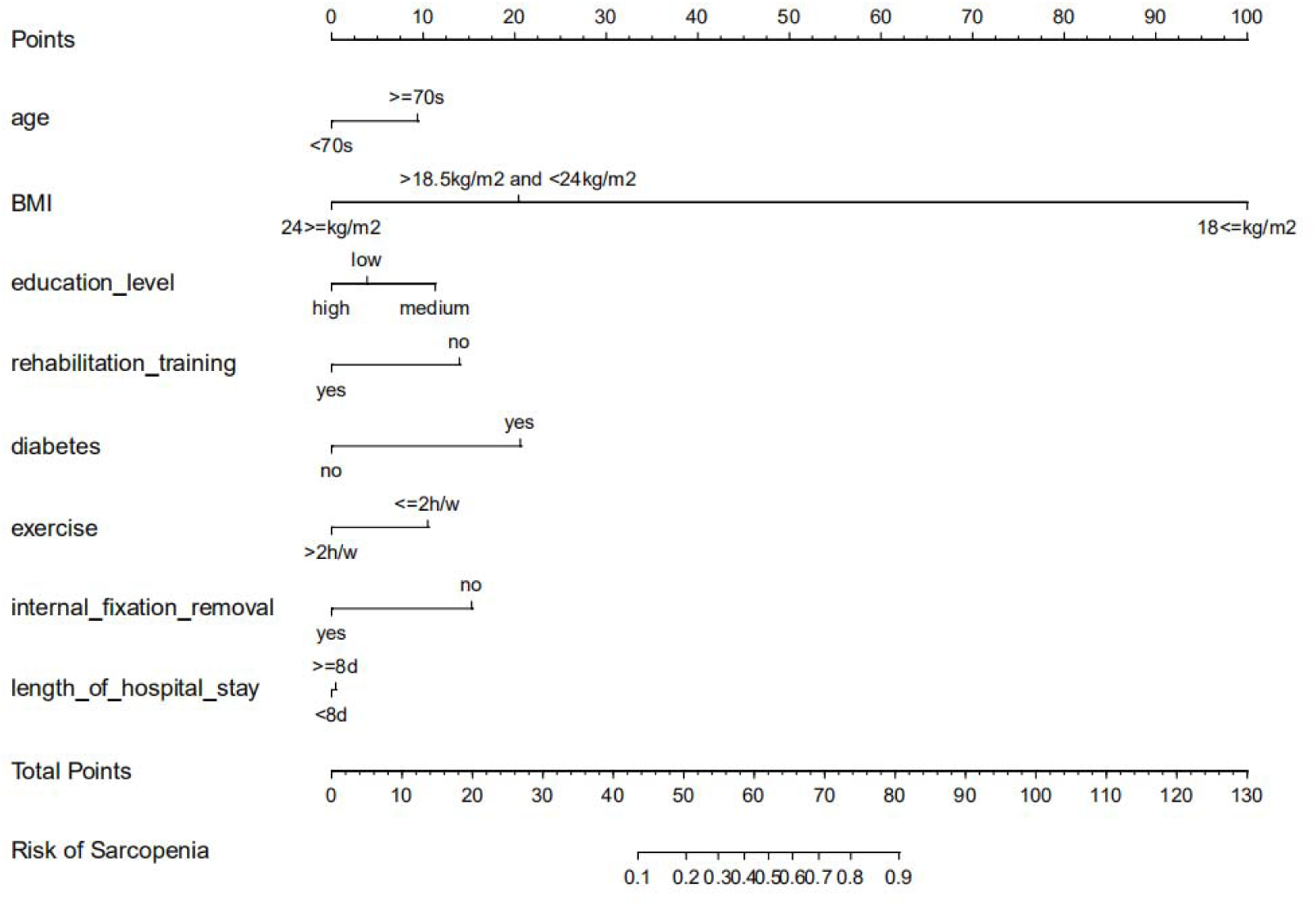
A nomogram model to predict the risk of sarcopenia after patella fracture. Note: 8 factors including age, BMI, education, postoperative rehabilitation training, diabetes, postoperative hospitalization, removal of internal fixation and regular outdoor exercise were included.

### Evaluation the nomogram

The calibration curve of the nomogram showed good agreement (Fig. 3a), which suggested that this model could be used to predict the risk of sarcopenia in patients after patellar fracture. In addition, the AUC of this prediction model was 0.8446115 (receiver operating curve shown in Figure 3b), and the C-index was 0.909 (95% confidence interval: 0.81–1.00). The C-index was determined to be 0.828 for the sample validation experiment in this study. This proved that this model was very efficient in terms of predictive power.

**Figure 3.**
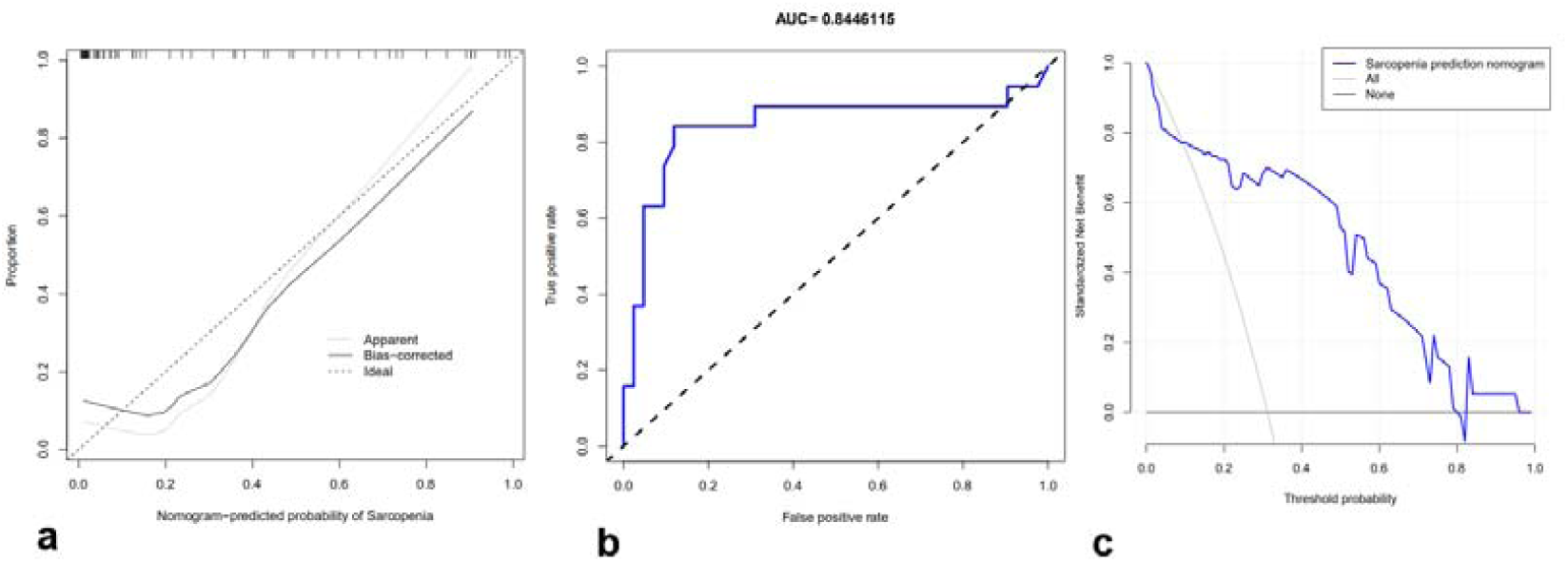
a. A calibration curve for the predictive model of muscle reduction in cases of patella fracture. Note: The x-axis is the risk of sarcopenia. The y-axis represents the actual incidence of sarcopenia. The diagonal dashed line represents a perfect prediction of an ideal model. The solid line indicates the predictive power of this predictive model, and the more it fits with the dotted line, the better the predictive ability. b. The area under the curve (AUC) of the muscle reduction prediction model indicates the probability of accurately predicting whether or not the patient has sarcopenia in a randomly selected case. This model has a good predictive power with AUC=0.8446115. c. The decision curve of the model for predicting muscle loss. The y-axis does not have the possibility of sarcopenia. The dotted line indicates the prevalence of muscle reduction predicted by the model. The solid line indicates that all patients have sarcopenia. The solid blue line indicates the likelihood of sarcopenia after a full interventional prophylaxis with this predictive model. The decision curve shows that if the risk of sarcopenia is between 8.3% and 80.0%, this predictive model has good clinical applicability. In this case, using the preventive program in this study can prevent the risk of sarcopenia better than preventive measures.

### Clinical application

The decision curve analysis of the nodule prediction model for postoperative sarcopenia in cases of patellar fractures is shown in Figure 4. The decision curve shows that for predicted probabilities of 8.3–80%, clinical interventions based on the nomogram prediction model can achieve a net rate of return similar to that of the predicted outcome. This implies that the current prediction model can better guide clinical practice and facilitate early planning and implementation of clinical intervention to reduce the risk of sarcopenia, thus reducing the overall risk of muscle loss in the population.

## Discussion

With the development and availability of R software, the nomogram prediction model is now being widely used in prognostic evaluation and assessment of treatment efficacy. Additionally, the nomogram prediction model has well-defined quantitative indicators, which facilities accurate evaluation of treatment outcome and prognosis.^31^ In this paper, we describe, for the first time, the use of a nomogram model for the assessment of the risk of post-fracture sarcopenia.

In this study, we developed and validated a new tool for predicting postoperative sarcopenia in patients with patellar fractures by using only eight easily available variables. Our preliminary tests reveal that this model can serve as a relatively accurate predictor of whether or not muscle loss occurs in patients who undergo surgery for patellar fractures. Internal sampling showed that the model has strong predictive ability, while high values of C-index and AUC suggest that this model can be widely used for the prediction of postoperative sarcopenia in patients, with good accuracy.^31^

In our study population, 31% of the patients developed sarcopenia. Analysis of the risk factors for sarcopenia included the following factors: age, BMI, education level, postoperative rehabilitation training, diabetes, postoperative hospital stay, removal of internal fixation, and regular outdoor exercise. Our predictive model showed that low BMI, diabetes, lack of rehabilitation after surgery, lack of regular exercise and removal of internal fixation may be the key individual factors that lead to postoperative muscle loss.

As shown in several other studies, a decrease in BMI is often indicative of a decrease in the muscle strength of the limbs,^32,33^ and patients with sarcopenia generally have relative low BMI.^34^ Therefore, a decline in BMI is an important manifestation of sarcopenia in patients undergoing surgery after fracture. In addition, studies have also shown that very low weight is closely related to the occurrence of sarcopenia.^35^ For the elderly, while obese body types can induce and worsen many diseases, low body weight is also associated with poor health and is an important risk factor for death.^36^ High BMI can be considered as protective against the development of sarcopenia in the elderly.^35^ Therefore, maintaining a high BMI through intake of adequate nutrients can prevent the occurrence of postoperative sarcopenia in elderly fracture patients. ^37^

More than 422 million people worldwide have been estimated to have diabetes,^38^ and sarcopenia has been reported as a complication of diabetes.^39,40^ Our study confirmed that diabetes was an important risk factor for muscle reduction in patients with patellar fractures. The incidence of sarcopenia has also been reported to be significantly high in patients with poor glycemic control, and sarcopenia in turn leads to the worsening of diabetes.^41^ This is because in the presence of type 2 diabetes, insulin resistance results in autophagy, degradation of muscle protein (ubiquitin–proteasome proteolytic pathway), and mitochondrial dysfunction; these processes eventually lead to the loss of muscle mass and muscle strength. Muscle mass and loss of muscle strength, in turn, lead to a potential increase in insulin resistance; additionally, further progression of mitochondrial dysfunction can aggravate insulin resistance, ultimately compromising diabetes control and then stimulating pathways leading to further muscle loss.^42-49^ This vicious circle between muscle reduction and diabetes can be broken by rehabilitation exercise, which is known to have a beneficial effect on bone metabolism.^50^ Further, studies have shown that aerobic exercise with appropriate resistance training is an economic and effective intervention and that it is helpful for effectively controlling diabetes and improving the condition of elderly diabetic patients.^51^ These findings are consistent with the results of our study, where our predictive model demonstrated that postoperative rehabilitation exercise and regular physical exercise can prevent the development of sarcopenia.^52^

Interestingly, we found that plate removal is a particularly important predictor of functional recovery of the knee joint after surgery. Retention of internal fixation has many risks, including screw fracture, postoperative fracture, important tissue damage, major bleeding, intraoperative fracture, infection, poor wound healing, etc.^53^ However, with thorough preoperative preparation, these complications were avoided in our patients. Additionally, some patients may develop limited joint mobility due to the retention of internal fixation. Retention of internal fixation also hinders the removal of heterotopic bone and soft tissue release, so it must be taken out for complete release of the tissues.^54-56^

Our prediction model also highlights other risk factors for the development of sarcopenia. Baumgartner et al have shown that the incidence of sarcopenia in patients aged between 65 and 70 years ranges at 13–24%.^57^ Further, studies have also shown that sarcopenia may occur in as many as half of elderly subjects aged ≥80 years.^58,59^ In addition, perhaps higher-educated patients may have greater awareness regarding health issues rather than patients with low education levels and may tend to have better prognosis than others.^60^ Our study also suggests that prolonged hospital stay may be a poor prognostic factor for fracture patients; this is consistent with the findings of previous studies that have shown that prolonged hospitalization affects patients’ daily activities.^61^

The predictive tool developed by us offers several benefits: (1) Using a predictive tool for postoperative muscle reduction in patients with patellar fractures enables individualized risk prediction, which can indirectly improve the overall patient outcomes.^62^ (2) Our model can serve as an effective sarcopenia risk prediction tool, which would enable clinicians to identify patients at a high risk of developing sarcopenia ad take appropriate measures to prevent the same. (3) Further, our assessment tool can provide a theoretical background to guide further clinical research on sarcopenia. For example, our nomogram prediction model can be used to facilitate the selection of patients with high risk of sarcopenia in clinical trials. (4) Furthermore, simple early interventions, such as the engagement of medical reminders and family support, will help reduce the risk of postoperative sarcopenia in high-risk patients.^63^ In summary, patients undergoing post-fracture surgery should be recommended treatment for diabetes, post-operative rehabilitation, regular physical activity, and education regarding postoperative management in order to reduce the risk of sarcopenia.^64^

Therefore, an accurate prognostic assessment tool would enable physicians to evaluate the risk of sarcopenia in a timely manner and accordingly make necessary interventions to improve the prognosis of patients, while also eliminating the wasteful use of medical resources caused by false-positive results. Although it is difficult to completely eliminate the occurrence of sarcopenia, it may be certainly possible to reduce its risk through reasonable evaluation and cost-effective interventions. This would substantially improve the outcomes in patients undergoing post-fracture surgery and therefore our findings have important clinical significance.

Our current study also has a few limitations. First, our sample population contains fewer males than females due to the higher risk of patellar fractures in the latter. Thus, this cohort is not representative of all patients with patellar fractures in China. Second, for the risk factor analysis in this study, we did not include all the risk factors for muscle reduction after patellar fractures. Third, although our prediction model has good internal sampling, the inclusion of stable external data is still necessary for further validation. Fourth, the data used in this study were collected from the patients’ clinical history and long-term follow-up data; although we take all measures to ensure accuracy of the data as much as possible, because of the high patient number, it may not be entirely possible to rule out missing or incorrect information.

## Conclusion

In this paper, we described a new predictive model that helps clinicians assess the risk of sarcopenia with good accuracy, in elderly patients with patellar fractures. Our findings indicated that the most common risk factors for muscle reduction in patients after fractures include advanced age, low BMI, diabetes, and long postoperative hospital stay. On the other hand, higher education level, postoperative rehabilitation training, timely removal of internal fixation, and regular outdoor exercise were identified as factors protecting against muscle loss. We believe that after further validation through statistical testing and trials on random population samples, this model can be economically and effectively used for predicting the of postoperative muscle reduction in elderly patients with patellar fractures. Accurate estimation of individual risks can help clinicians and patients adopt appropriate modifications in lifestyle and medical interventions. Further research is warranted to determine whether change in interventions based on this nomogram prediction model can effectively reduce the risk of sarcopenia and improve patient outcomes.

## Acknowledgements

This work was supported by National Natural Science Foundation of China grants (81371963) and Project of Shanghai Jiaotong University (601N16025). Thanks for the anonymous reviewers for their valuable comments and suggestions that helped improve the quality of our manuscript

## Competing interests

The authors declare that they have no competing interests.

## Availability of data and materials

Please contact author for data requests.

## Authors’ contributions

Yisheng Chen and Jie Tao* designed the study. Yisheng Chen wrote the manuscript. Jie Tao, Yanxian Cai revised and polished the manuscript. Jie Tao, Yisheng Chen, Xueran Kang and Zhou ZH performed the experiments and statistical analysis of the data. Zhou ZH, Chenting Ying and Yunpeng Zhang conducted some experiments. Xueran Kang, Qi Xin, Yanxian Cai and Jiang Yang provided some advice and collected some relevant literature. All authors read and approved the final manuscript.

## Ethics approval and consent to participate

This study was approved by the Institutional Ethics Review Board of Shanghai General Hospital, Shanghai Jiao Tong University, Shanghai, China.

